# A Novel In Situ Activity Assay for Lysyl Oxidases

**DOI:** 10.1101/2021.04.30.442175

**Authors:** Huilei Wang, Alan Poe, Lydia Pak, Sandeep Jandu, Kavitha Nandakumar, Jochen Steppan, Reik Löser, Lakshmi Santhanam

**Author notes:** To whom correspondence should be addressed: Lakshmi Santhanam, Ph.D., Johns Hopkins University SOM, 720 Rutland Avenue, Ross 1150, Baltimore, MD 21205, USA.

## Abstract

The lysyl oxidase family of enzymes (LOXs) catalyze oxidative deamination of lysine side chains on collagen and elastin to initialize cross-linking that is essential for the formation of the extracellular matrix (ECM). Elevated expression of LOXs is highly associated with diverse disease processes. To date, the inability to detect total LOX catalytic function in situ has limited the ability to fully elucidate the role of LOXs in pathobiological mechanisms. Using LOXL2 as a representative member of the LOX family, we developed an in situ activity assay by utilizing the strong reaction between hydrazide and aldehyde to label the LOX-catalyzed allysine (-CHO) residues with biotin-hydrazide. The biotinylated ECM proteins are then labeled via biotin-streptavidin interaction and detected by fluorescence microscopy. This assay detects the total LOX activity in situ for both overexpressed and endogenous LOXs in cells and tissue samples and can be used for studies of LOXs as therapeutic targets.

## Introduction

The lysyl oxidase (LOX) family of enzymes play a critical role in the formation, maturation, and remodeling of extracellular matrix (ECM)^1^. The LOX family is composed of the prototypical LOX and four LOX-like proteins (LOXL1-4), all of which have a highly conserved C-terminal amino acid sequence (the LOX domain) that is required for amine oxidase catalytic activity^2,3^. Both the prototypical LOX and LOXL2 are critical for embryonic development, and germline deletion results in high embryonic or perinatal lethality^4,5^. Conversely, the sustained activation of LOX and LOXL2 causes significant ECM deposition, mechanical stiffening, and fibrosis^6,7^. Upregulation of LOX and LOXL2 has been implicated in several pathologies, including cancer, cardiac fibrosis, idiopathic pulmonary fibrosis, vascular stiffening in aging, and pulmonary hypertension^8–14^. Recent advances have placed LOXL2 as an appealing target in these diseases and generated substantial interest in the development of specific and selective inhibitors^15,16^.

Although a number of studies have shown protein and mRNA expression of lysyl oxidases (LOXs) to be significantly elevated in pathological specimens when compared to that of healthy controls^17–19^, the measurement of enzymatic activity of LOXs in diseased vs. healthy tissues has remained challenging. LOXs catalyze the oxidative deamination of lysine and hydroxylysine side chains of collagen and elastin precursors to produce highly reactive semialdehydes called allysines. The allysines spontaneously condense to form covalent bonds that crosslink collagen and elastin^20^. To date, studies have relied on two techniques to detect LOXs activity in tissue specimens. The first method uses the detection of hydrogen peroxide (H_2_O_2_) released as a byproduct during the LOXs catalytic cycle (**Fig. 1**) as a surrogate measure of LOX activity in solubilized samples^21^. Released H_2_O_2_ is detected using Amplex red in horseradish peroxidase (HRP)-coupled reactions. This method suffers from three key drawbacks when used to interrogate LOX activity in cells and tissue: (1) The primary interest is often to measure the activity of LOXs in the ECM, and the harsh conditions (e.g., SDS or urea) required to solubilize and release LOXs from the ECM of tissue samples into solution can cause loss of activity^1,22^; in addition, SDS inactivates HRP^23^ and urea denatures HRP^24,25^, which render the assay unreliable; (2) The approach requires the pooling of samples to obtain an observable signal, a significant pitfall when examining small, precious specimens; and (3) H_2_O_2_ can be produced from other enzymatic sources in cells and tissue^26^, generating significant noise in the assay; this is specifically a concern in fibrosis, wherein ROS pathways are often upregulated^27^. The second approach to detect LOXL2 activity specifically measures collagen IV 7S dodecamer crosslinks in the ECM by Western blotting^28–30^. This method is more robust and is able to produce low-background results for LOXL2 crosslinking of the COLIV 7S subunit and has been successfully used in various applications^28–30^. However, this assay also poses the following challenges: 1) the upstream steps to prepare the ECM specimens for Western blotting in this assay are involved, time consuming, and nuanced. Briefly, the detergent-insoluble ECM is first digested with collagenase. Then, collagenase resistant 7S dodecamers are extracted and purified by ion exchange and gel filtration chromatography; 2) this method is not suitable for in situ application and does not yield information on the specific regions within a tissue where LOXL2 activity is upregulated in disease; 3) Prior studies have shown that LOXL2 can oxidize other matrix proteins including type I collagen^31^ and tropoelastin^32^. Therefore, this assay cannot fully capture overall LOXL2 activity in the matrix, nor does it yield overall LOXs activity. Thus, a sensitive and selective assay that reliably detects the activity of LOX isoforms in situ and can be adapted for specific LOX isoforms in the ECM is still needed.

**Figure 1.**
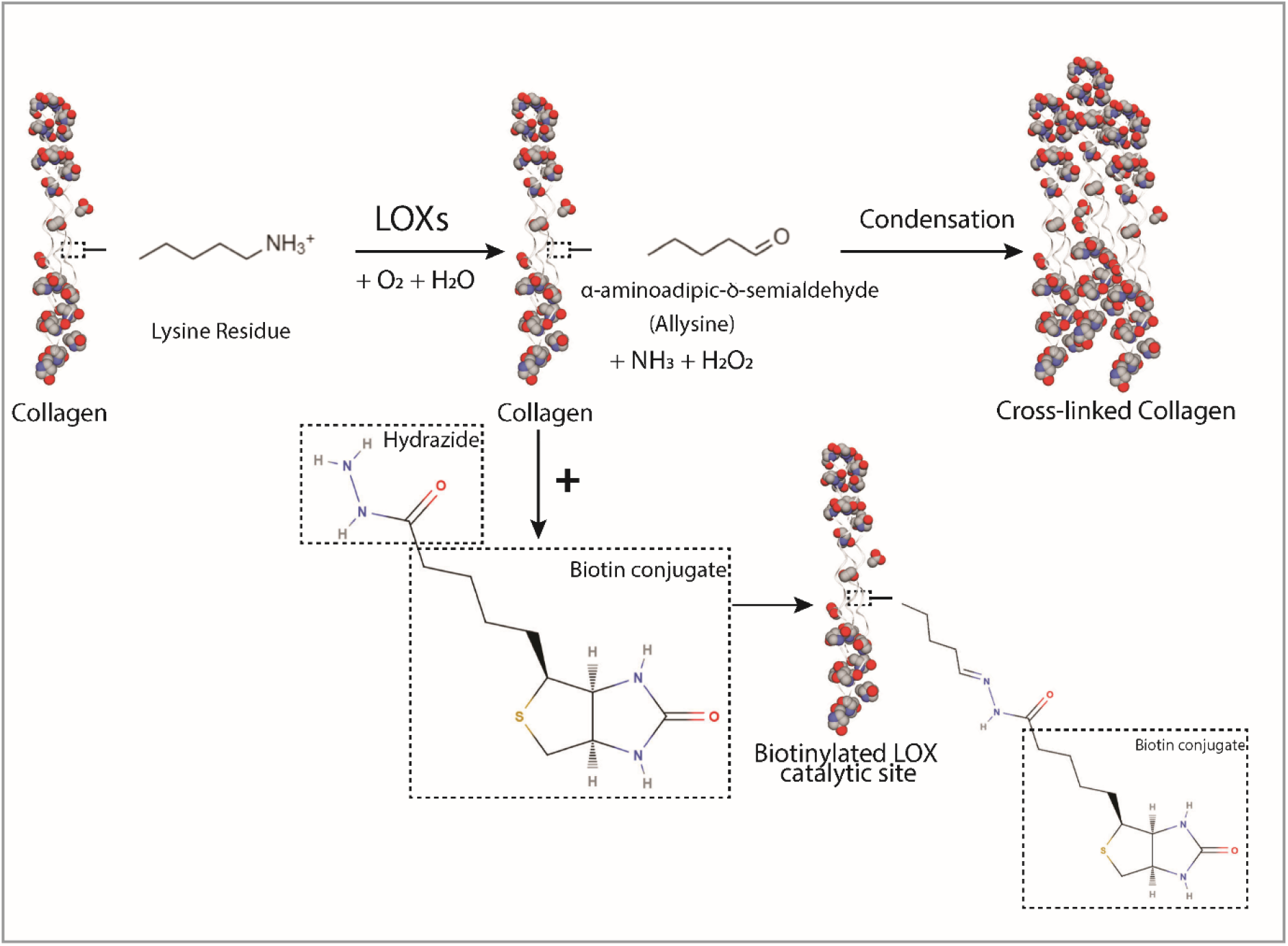
Oxidative deamination catalyzed by lysyl oxidases (LOXs) leads to collagen crosslinking by spontaneous condensation. Intermediate semialdehyde and the imine and aldol products of the crosslinking reaction can be labeled by biotinylated hydrazide. LOXs oxidize lysine and hydroxylysine side chains on collagen and elastin to highly reactive semialdehydes called allysines, while producing hydrogen peroxide (H_2_O_2_) and ammonia (NH_3_) as byproducts. Allysines then spontaneously condense to form cross-linkages. In our approach to detect LOXL2 activity, adding biotin-hydrazide (BHZ) results in biotinylation of LOXL2-catalyzed peptidyl lysines.

In this study, we exploited the reactivity of hydrazides with imines and carbonyls to develop a new in situ LOXs activity assay^33,34^. A prior study has shown that LOXL2-dependent generation of aldehydes on PDGFRβ can be labeled with biotin hydrazide and detected in a pulldown assay with avidin-coupled affinity resin^35^. We hypothesized that this strategy can be generalized to detect the overall enzymatic activity of LOXs. Specifically, the imine and allysine intermediates generated by lysyl oxidase activity are labeled directly with biotin-hydrazide (BHZ; **Fig. 1**). This reagent reacts with the carbonyl group that is present in both the allysine residues formed as immediate products of LOXs-catalyzed deamination and in the products of the subsequent spontaneous crosslinking reaction of the aldol condensation type. Furthermore, it can be assumed that BHZ also reacts with the predominantly formed Schiff base-type crosslinking products^36^, since it is known that aldimines react analogously to aldehydes with hydrazine derivatives to form hydrazones^37,38^. Therefore, a stoichiometric excess of BHZ should allow for the favorable detection of the allysine residues and imines derived thereof formed by the enzymatic activity of LOXs. The biotin-streptavidin interaction is then harnessed for fluorescence-based detection of BHZ-labeled proteins that are visualized by epifluorescence or confocal microscopy. In this study, we used LOXL2 as a representative member of the LOX family for proof of concept and validation assays because LOXL2 has recently garnered significant interest as a target in fibrosis and vascular aging^12,15,16^.

## Results

### Proof of principle

For proof of principle, we used gain-of-function and loss-of-function approaches to identify if increased or decreased LOXL2 protein expression and activity directly lead to increased or decreased BHZ incorporation, respectively. First, in gain-of-function experiments, we examined whether overexpression of secreted, catalytically active LOXL2 increases BHZ incorporation in vascular smooth muscle cells (VSMCs). Viral transduction of catalytically active LOXL2 in A7r5 rat aortic VSMC resulted in a significant increase in BHZ (100 μM; 24 h) incorporation in the extracellular space when compared with control (baseline) A7r5 cells (**Fig. 2A**). Areas with high biotinylation signal showed fibrous structures, similar to structures of cross-linked ECM proteins. To further verify that the catalytic activity of LOXL2 was responsible for the increase in BHZ signal, we used the following negative controls: 1) the overexpression of catalytically inactive H626/628Q LOXL2 double mutant (LOXL2-DM)^39,40^, 2) inhibition of LOXL2 specific activity using the small molecule inhibitor PAT-1251 (10 μM) in cells overexpressing catalytically active LOXL2^41^, and 3) inhibition of total LOX activity using the non-selective small molecule LOX inhibitor β-Aminopropionitrile (BAPN; 10 μM) and LOXL2-specific inhibitor PAT-1251^41,42^ in cells overexpressing catalytically active LOXL2. In all these three conditions, BHZ incorporation was low, showing no difference from control cells (**Fig. 2Ai**). Image analysis to quantify BHZ incorporation showed a 2-fold increase in BHZ signal with overexpression of wild-type LOXL2, but not with overexpression of inactive LOXL2-DM, or in the presence of inhibitors (bar graph, **Fig 2Aii**). LOXL2 and LOXL2-DM overexpression by adenoviral transduction was verified by Western blotting. LOX, LOXL1, and LOXL3 expression were not altered by LOXL2 or LOXL2DM overexpression (**Fig 2Aiii**).

**Figure 2.**
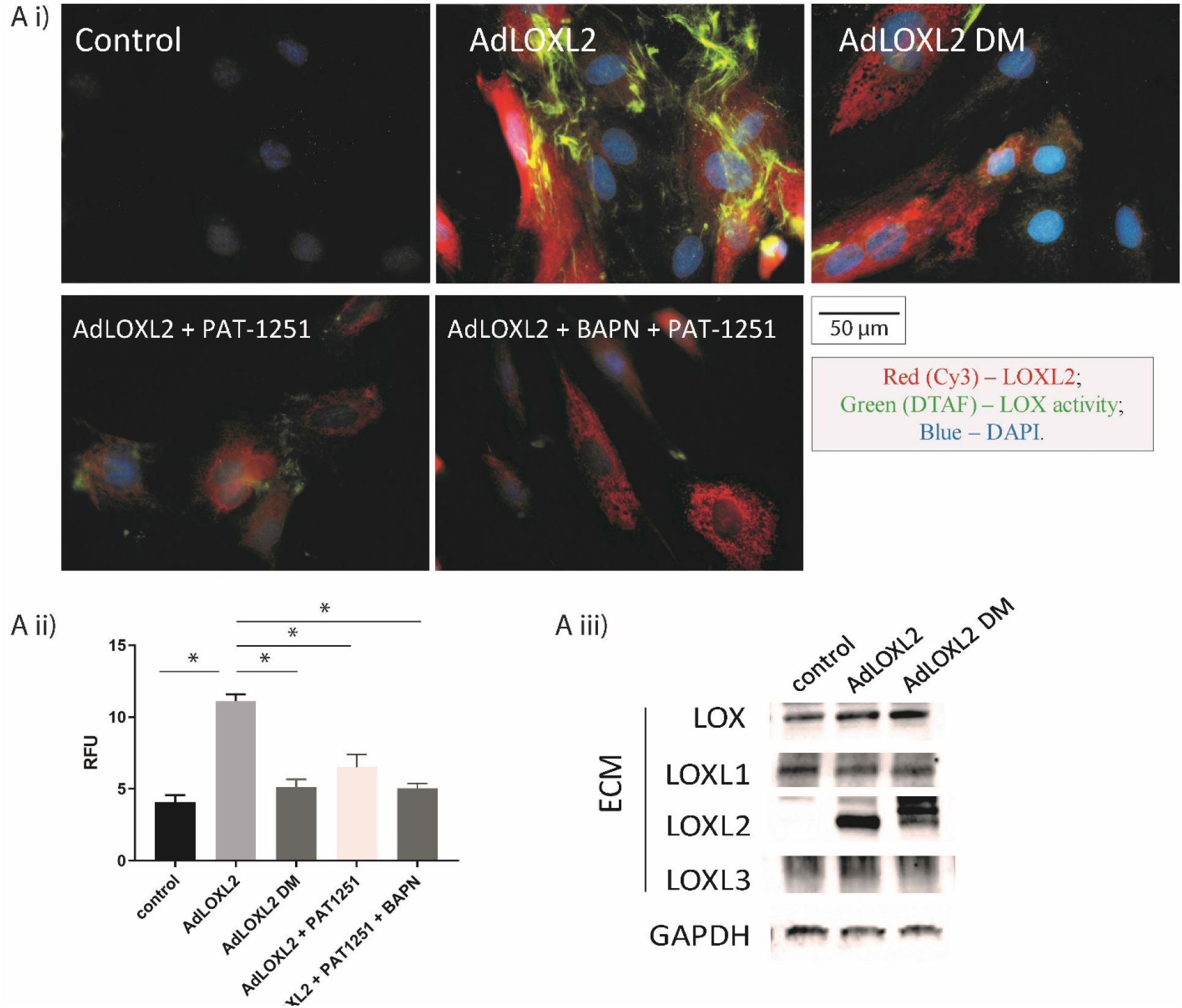
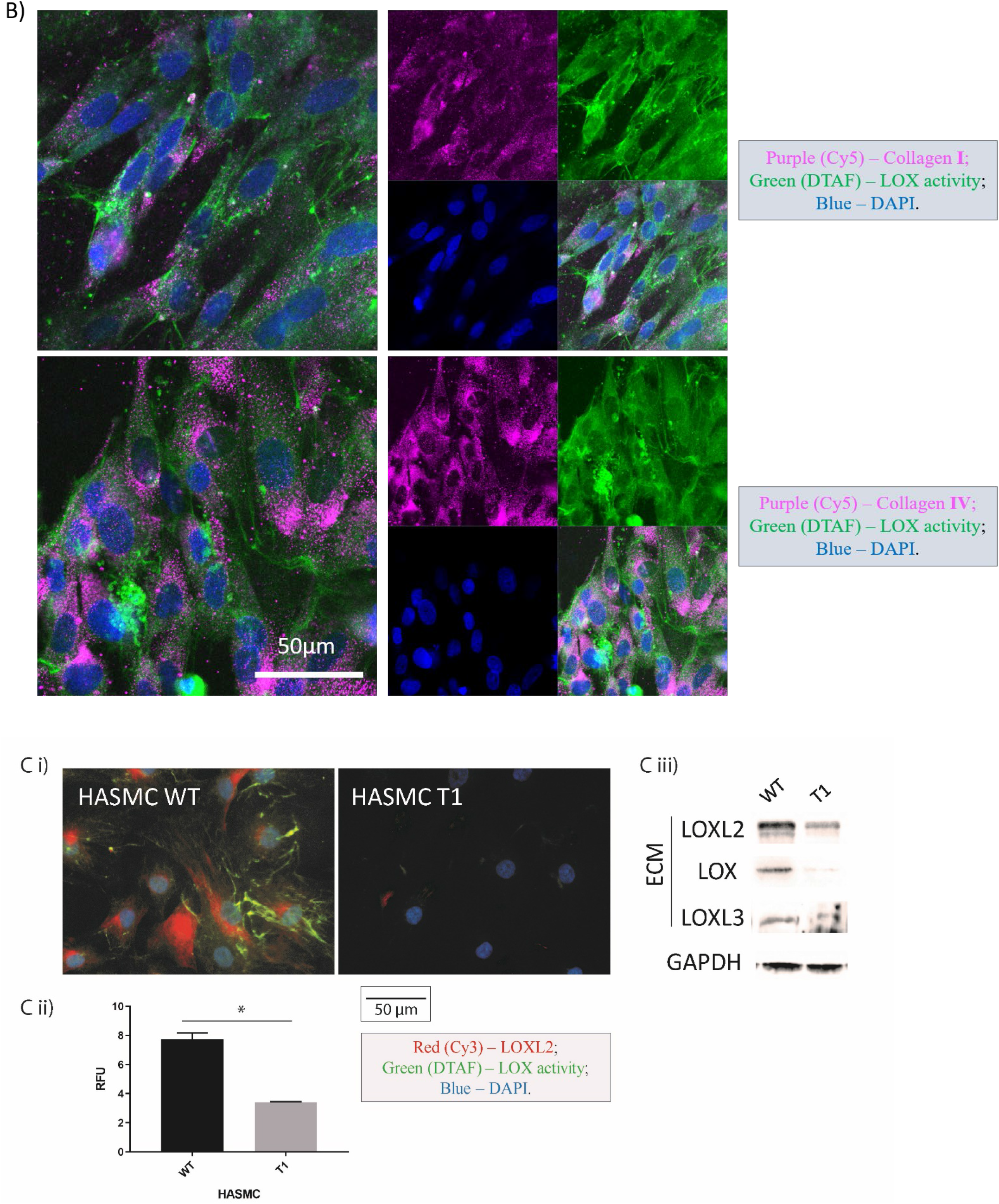
Proof of concept for the in situ LOXs activity assay. Cells were incubated with 100 μM biotin-hydrazide (BHZ) for 24 h and then fixed and co-stained for LOXL2 (immunostaining; red), biotinylation (DTAF-streptavidin; green) corresponding to LOXs activity, and nuclei (DAPI; blue). **A)** Representative confocal microscopy images of i) control A7r5s cells, ii) wild type LOXL2 overexpression iii) catalytically inactive LOXL2-DM overexpression, iv) LOXL2 overexpression + LOXL2-specific inhibitor PAT-1251 (10 μM), v) LOXL2 overexpression + inhibitors BAPN (10 μM) + PAT-1251 (10 μM). (n = 12; Scale bar = 50 μm). Activity signal in each IF image was converted to mean gray value shown in bar graph (mean±SEM; *P<0.05 by student’s t-test). Representative western blotting images compare the expression of LOXs in the ECM of control A7r5 cells and A7r5 cells with wild type LOXL2 or LOXL2-DM overexpression. **B)** Representative confocal immunofluorescence microscopy images of A7r5 cells with LOXL2 overexpression. BHZ incorporation (green) was co-stained with collagen I or collagen IV (purple), and nuclei (blue). (n=5; Scale bar = 50 μm.) **C)** Representative confocal microscopy images of LOXs activity assay performed in wild-type human aortic smooth muscle cells (HASMC) and CRISPR-Cas9 mediated LOXL2 gene knockout HASMC T1 cells (Ci; Scale bar = 50 μm). Bar graph shows measured activity signal (n=24; mean±SEM; *P<0.05 by student’s t-test) (Cii). Representative western blotting images compare the expression of LOXs in the ECM of wild type HASMC cells and HASMC T1 cells (Ciii).

Next, we determined if the BHZ signal noted in the gain-of-function experiment colocalizes with type I and type IV collagens, which are the major extracellular protein substrates of LOXs by co-immunofluorescence staining. LOXL2-overexpressing A7r5 cells showed significant overlap of BHZ incorporation and endogenous collagen I and IV deposited in the ECM (**Fig 2B**), which indicates that the BHZ-derived signal originates from the LOX-catalyzed conversion of lysine-containing proteins.

Next, we used a loss-of-function approach as a second line of evidence (**Fig 2C**). We compared human aortic smooth muscle cells (HASMCs) with and without CRISPR-Cas9 editing of the LOXL2 gene (HASMC T1)^12^ to test whether LOXL2 depletion correlates with loss of BHZ incorporation signal and to determine the specificity of the assay. Western blotting was used to confirm that LOX2 was indeed depleted in the HASMC T1 cells. Interestingly, in addition, the abundance of LOX and LOXL3 was also reduced in the HASMC T1 cells (**Fig 2Ciii**), showing a significant decrease in total LOXs in the HASMC T1 cells. BHZ incorporation in the ECM was attenuated by 66% in the ECM of LOXL2-depleted HASMC T1 cells exhibited when compared to wild-type (WT) HASMCs (**Fig 2Ci,ii**). This result confirmed that the 1) the assay detects endogenous LOXs activity in cells, 2) extracellular incorporation of BHZ corresponds to the presence of total activity of LOXs in the extracellular space, and 3) LOXL2 depletion results in a significant loss of total LOXs activity in HASMC.

### Assay Specificity

To further explore the specificity of our assay towards LOXs, we examined three potential sources of nonspecific signal: 1) other amine oxidases, 2) small molecule aldehydes generated from reactive oxygen species (ROS), and 3) BHZ – LTQ interaction.

As other LOXs or amine oxidase, copper containing 3 (AOC3) are produced by VSMCs and could be responsible for the increase in BHZ signal^43,44^, A7r5 cells were subjected to immunostaining to measure changes in expression levels of these proteins when LOXL2 or LOXL2-DM is overexpressed (**Fig S1**). Results showed that the expression of other LOXs and AOC3 were unaffected by AdLOXL2 and AdLOXL2-DM viral transduction. Furthermore, at 10 μM concentration, BAPN would exclusively inhibit lysyl oxidases (IC_50_(LOX)=5 μM), whereas SSAO would not be affected at this concentration (IC_50_=4 mM)^45^. Similarly, PAT-1251 would inhibit both LOXL2 and LOXL3 at 10 μM (IC_50_(LOXL2)=0.87 μM, IC_50_(LOXL3)=1.38 μM), but inhibit less than 10% inhibition of hDAO and hMAO^41^. The efficient suppression of BHZ incorporation under these conditions suggests that the contribution of other amine oxidases in this assay can be ruled out.

**Figure S1:**
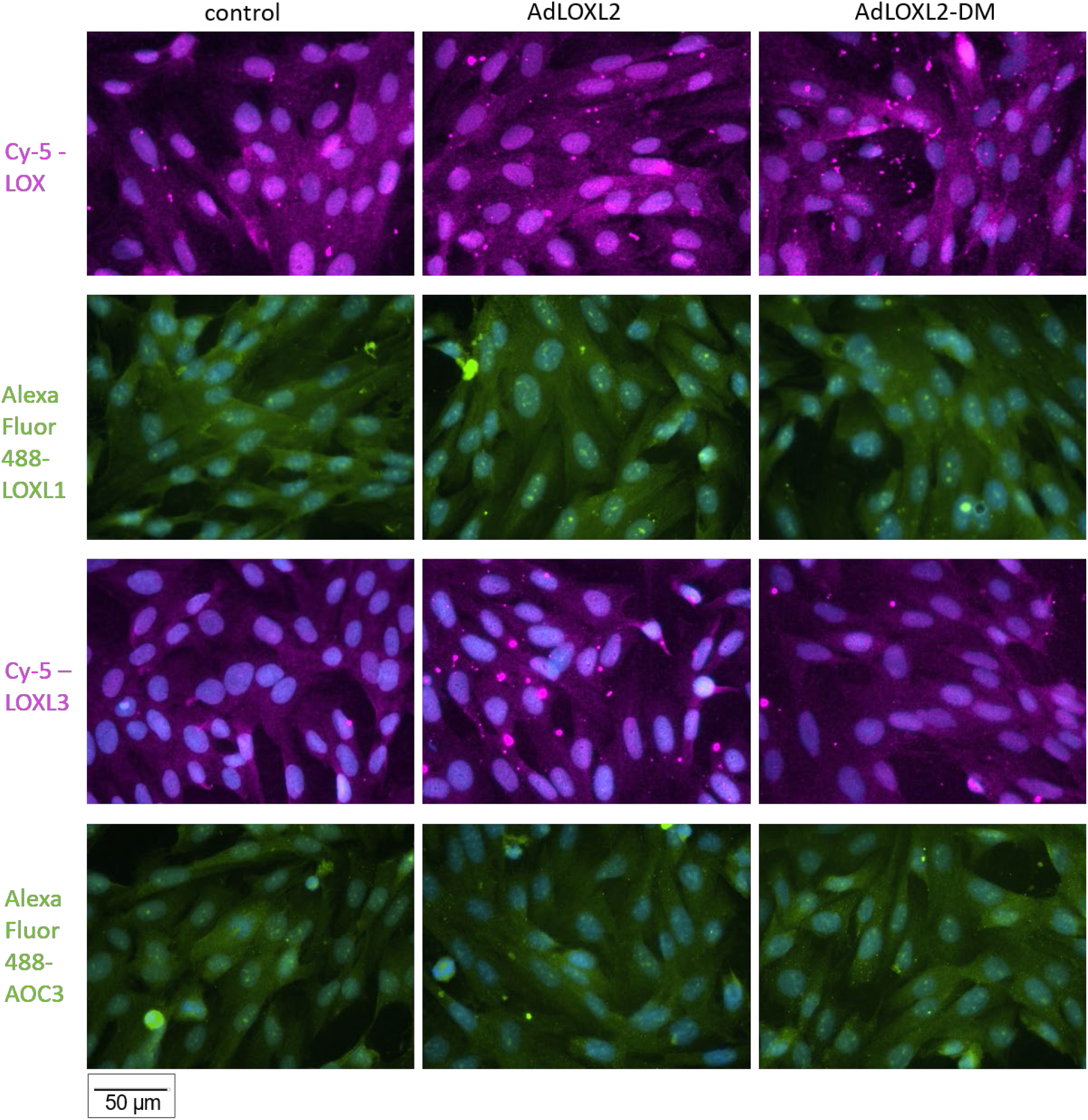
Representative immunofluorescence microscopy images of A7r5 cells transduced with AdLOXL2 or AdLOXL2-DM. Cells overexpressing LOXL2 or LOXL2DM were stained for LOX, LOXL1, LOXL3 and AOC3 separately, and co-stained with DAPI (blue). No significant increase in expression was seen for these proteins in AdLOXL2 and AdLOXL2-DM cells.

Small molecule aldehydes can also arise from the oxidation of unsaturated fatty acids (linoleic acids in the case of C9-aldehydes) in membrane lipids from reactive oxygen species generated under oxidative stress^46^. The application of vitamin E should largely attenuate lipid peroxidation, and therefore was used as a control in our assay to rule out the contribution of small molecule aldehydes generated from ROS^47^. Results (**Fig S2**) showed that the difference in BHZ incorporation was nonsignificant with the addition of Vitamin E (α-Tocopherol, 50μM). Taken together, these findings confirmed that the increase in BHZ signal observed in the LOXL2–overexpressing cells is due to LOXL2 catalytic activity.

**Figure S2:**
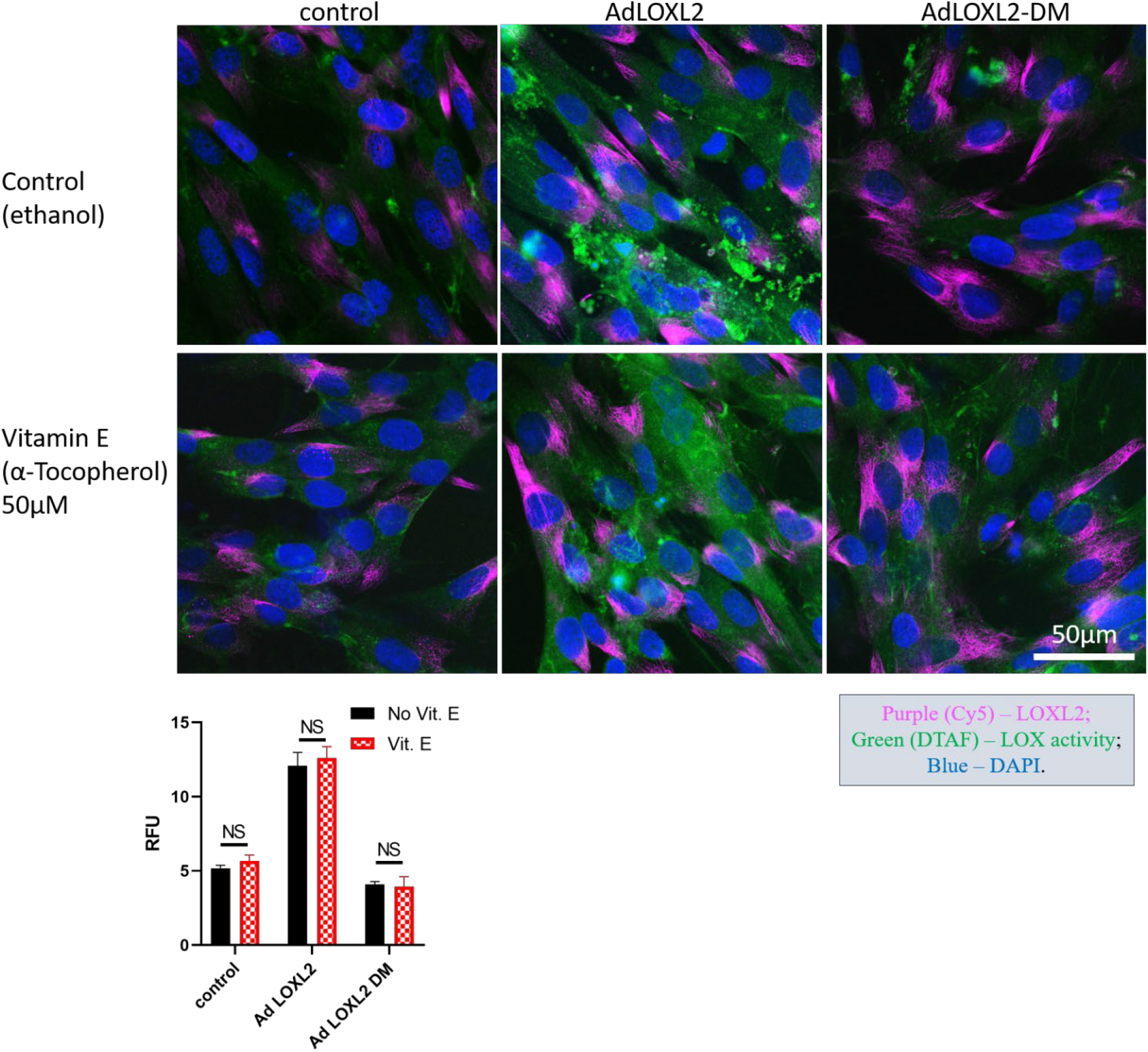
Representative confocal microscopy images of A7r5 cells transduced with AdLOXL2 or AdLOXL2-DM. Cells were incubated with 100 μM biotin-hydrazide (BHZ), with or without vitamin E (50 μM α-Tocopherol) for 24 h and then fixed and co-stained for LOXL2 (purple; immunofluorescent staining), biotinylation corresponding to LOXs activity (green; DTAF-streptavidin), and nuclei (blue; DAPI). Activity signal in each IF image was converted to mean gray value shown in bar graph (n=12). BHZ incorporation was unchanged in cells with the application of vitamin E. (n = 6; Scale bar = 50 μm. *P<0.05 by student’s t-test.)

The activity of lysyl oxidases is highly dependent on cofactor lysyl tyrosylquinone (LTQ)^48^. LTQ first reacts with the lysine ε-amino group to form a substrate-LTQ Schiff base adduct, which then undergoes abstraction of α-proton to form quinolaldimine. Hydrolysis of the adduct releases allysine^49^, which subsequently undergoes spontaneous reaction with another ε-amino group of lysine to form a Schiff base or spontaneously reacts with another allysine through aldol condensation^50^. Based on prior literature, biotin-hydrazide could detect the allysine products and Schiff base-type protein-protein crosslinks derived thereof on extracellular matrix proteins^38^, and potentially react with the carbonyl group on the LTQ and in turn inhibit LOX activity^33,51^. To determine whether the reaction of BHZ with LTQ generates signal in this assay, we overexpressed LOXL2 in A7r5 cells and incubated the conditioned cell culture medium, which contains high levels of catalytically active LOXL2 (Fig S3A) with increasing concentrations of BHZ for 2h or 24h, and enriched biotinylated proteins using streptavidin agarose beads. LOXL2-DM was used as a negative control in this assay, as the LTQ generation is a self-processing reaction that requires copper binding^52,53^ that is lacking in the LOXL2-DM^40^. Thus, any binding of BHZ to the LOXL2-DM protein would indicate the presence of other reactive functional groups in the LOXL2 protein that can generate noise in the assay. Increased LOXs activity was noted in the conditioned media from LOXL2 overexpressing cells, confirming the presence of LTQ cofactor whereas LOXL2-DM was catalytically inactive, as expected. Neither LOXL2 or LOXL2-DM was biotinylated at the various concentrations of BHZ (Fig. S3B), indicating no detectable reaction between BHZ and LTQ and no conjugation with LOXL2 protein. Thus, we conclude that the biotinylation noted in the extracellular space of A7r5 cells and HASMCs (Fig. 2) was due primarily to the reaction between BHZ and ECM protein modifications catalyzed by LOXL2 and not due to binding to the LTQ cofactor. This is in agreement with results obtained in a previous study, where 2,4-dinitrophenylhydrazine was used as reagent for antibody-based detection of allysine-containing cell-surface proteins^54^. Similar to BHZ, no reactivity of that hydrazine derivative towards other carbonyl-functionalized cellular constituents was noted.

**Figure S3.**
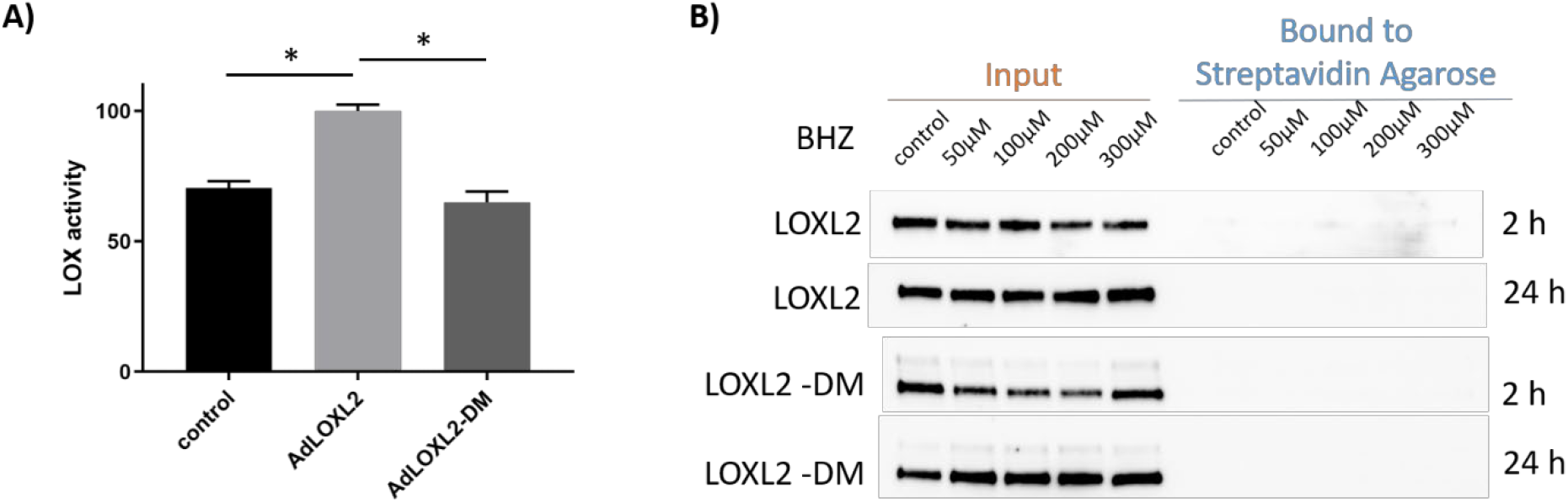
Biotin-hydrazide (BHZ) does not bind directly to LOXL2. **A)** H_2_O_2_-resorufin coupled LOX activity assay in cell culture medium from A7r5 cells with and without adenoviral-induced overexpression of LOXL2 or LOXL2-DM Activity was calculated as the initial rate of reaction calculated as the slope of fluorescence (relative fluorescence unit (RFU)) per unit time (1 h) normalized to the average signal of AdLOXL2 samples. (n=15; *P<0.05 by student’s t-test). **B)** Representative Western blot image comparing total and biotinylated LOXL2 and LOXL2DM after incubation with increasing concentrations of BHZ for 2h or 24h followed by enrichment with Streptavidin agarose beads. Western blots showed no meaningful levels of biotinylated-LOXL2.

### Optimization of experimental parameters

Next, to optimize the assay conditions, we investigated the effects of BHZ concentration (0-150 μM) and duration of labeling (24-72 h) on A7r5 cells that overexpressed LOXL2 via adenoviral delivery (Fig. 3). A linear BHZ concentration response was noted for 24 h and 48 h of incubation. The incubation of cells with 100 μM or 150 μM BHZ for 24 h yielded a robust signal within a reasonable timeframe for a cell-based assay. Although 50 μM yielded similar results at later time points, that concentration was suboptimal for this particular application because it increased the experimental duration and because cells are influenced by toxic effects at 48 h, likely due to the prolonged DMSO exposure.

**Figure 3.**
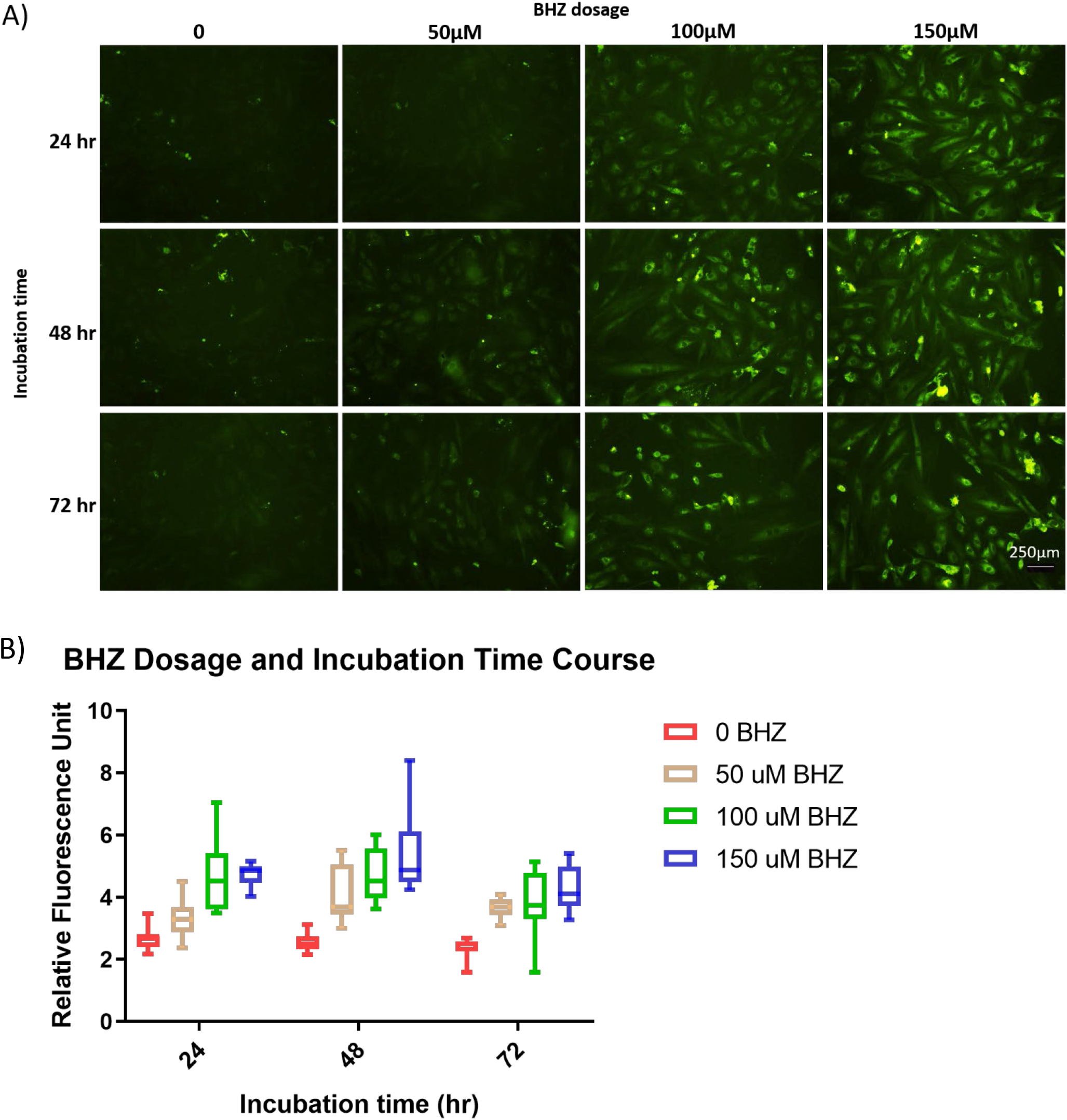
Optimization of LOXL2 activity assay parameters. **A)** A7r5 cells overexpressing LOXL2 were seeded on coverslips and incubated with different concentrations of biotin-hydrazide (BHZ; 0, 50, 100, or 150 μM) for 24, 48, or 72 h. After being fixed and stained for immunofluorescence with DTAF-streptavidin, cells were imaged with confocal microscopy in five independent coverslip locations per condition. (Scale bar = 250 μm). **B)** LOXs activity signal from each image was converted to grayscale and analyzed for mean gray value. (n = 10)

### Analytical application

Finally, we examined whether the assay can be used to detect increased LOXs activity in pathologic specimens. We have previously shown that aging is associated with significant upregulation of LOXL2 abundance and total LOXL2 activity in the aortic ECM^12^. Thus, we compared BHZ incorporation in freshly isolated aortas from young and old WT LOXL2+/+ and heterozygous LOXL2+/− littermate mice. Western blotting results confirmed that LOXL2 expression was highest in aortic tissues of old WT mice and lowest in those of young LOXL2+/− mice (Fig. 4A). Additionally, LOXL1 expression was significantly upregulated in aortic ECM of old mice compared to that of young mice. The overall LOXs content is the highest in old WT mice aortic ECM. Freshly isolated, live aortic specimens were incubated with BHZ (200 μM) for 24 h. After excess BHZ was removed by washing, the samples were fixed and co-stained with streptavidin DTAF to detect biotinylation signal and rabbit anti-LOXL2 followed by Alexa Fluor 568 Goat anti-Rabbit IgG. Consistent with Western blotting results, confocal immunofluorescence imaging of aortic rings showed that LOXL2 expression was higher in WT mice than in LOXL2+/− mice and higher in old mice than in young mice (Fig. 4B). Most important, total LOXs activity was strikingly higher in the aortic rings of old WT mice than in those from both young WT mice and old LOXL2+/− mice, as evidenced by a markedly higher level of BHZ incorporation. Aortic rings from young LOXL2+/− mice had the least LOXL2 activity when compared with rings from the other three groups. Control aortic rings incubated without BHZ had no biotinylation signal (Fig. 4C), confirming that levels of endogenous and nonspecific biotin are significantly below those produced by the LOXs activity assay, and that this approach of measuring LOXs activity has low background signal.

**Figure 4.**
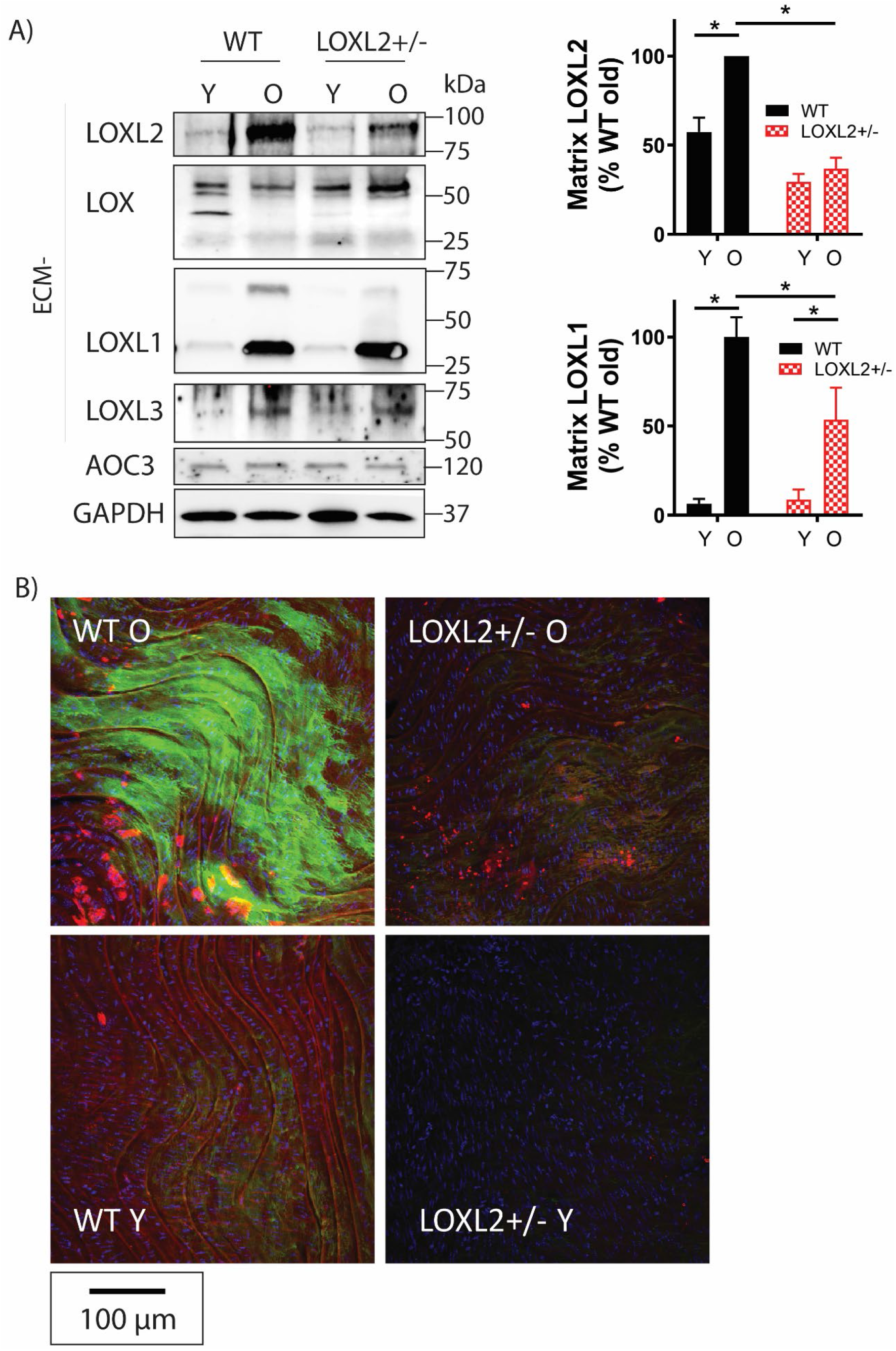
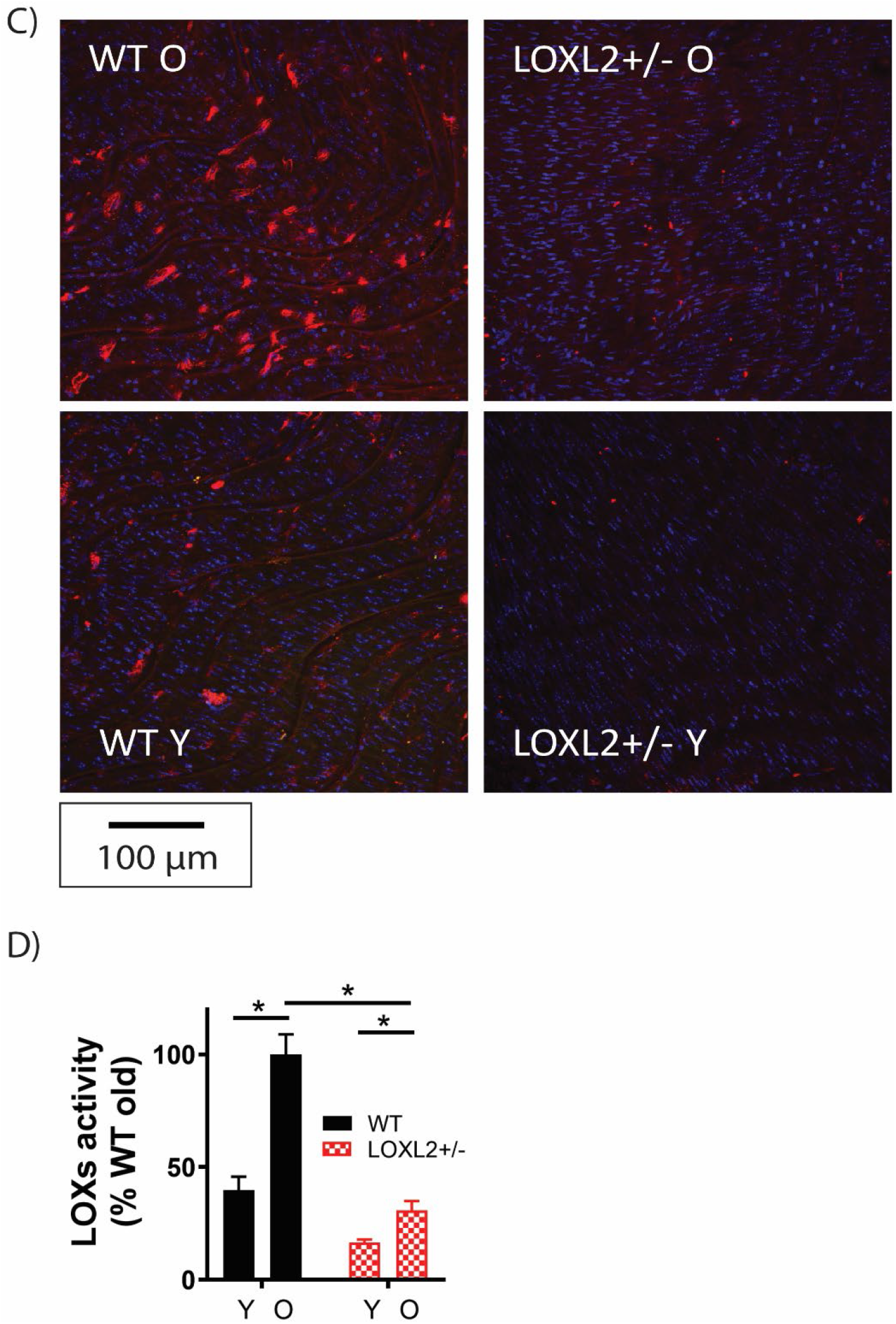
LOXL2 activity assay applied to aortic tissue samples. **A)** Representative Western blot images of LOXL2, LOX, LOXL1 and LOXL3 expression in aortic matrix of young (Y, < 3 months old) and old (O, > 18 months old) wild-type (WT) and LOXL2+/− littermate mice (n = 8). AOC3 is detected in the cytosolic fraction of aortic tissue homogenates but not in ECM and AOC3 expression level is similar across four groups. GAPDH was used as loading control. Bar graph shows the densitometry analysis of Western blots (mean ± SEM; n=6; *P<0.05 by 2-way ANOVA). **B)** Representative confocal Z-stack maximum projection immunofluorescence images show LOXL2 activity in aortic rings from young and old, WT and LOXL2+/− mice. Freshly isolated aortic rings were incubated with 200 μM biotin-hydrazide (BHZ) in DMEM + ITS for 24 h, then fixed and stained for LOXL2 (red), LOXs activity (green), and nuclei (blue). (n=6; Scale bar = 100 μm). **C)** Representative confocal images of control aortic rings incubated without BHZ. Samples were stained for LOXL2 (red), endogenous biotinylation (green), and nuclei (blue). (n = 6; Scale bar = 100 μm). **D)** Bar graph of LOXs activity calculated as mean gray value of the IF images (mean ± SEM; n=12, *P<0.05 by 2-way ANOVA). The overall LOXs activity in each group corresponds positively to the expression levels of LOXs in aortic ECM shown in A).

## Discussion

The LOX family enzymes have long been recognized to have an essential role in fibrogenesis. In recent years, LOX, LOXL1, and LOXL2 have emerged as potential therapeutic targets in a diverse set of diseases characterized by matrix remodeling and ECM deposition, including fibrosis, primary and metastatic cancers^8–14^. Our prior study showed that LOXL2 drives age-associated vascular stiffening by promoting ECM deposition and remodeling^12^. However, the specific role of LOXs in the ECM remodeling process remains incompletely understood, in part because of the lack of readily applicable, reliable assays to measure LOXs activity in the ECM. In this study, we used LOXL2 as the representative protein of the LOX family to develop and validate this new assay. However, the assay has the power to detect total in situ LOXs activity, which is key to overall matrix remodeling in a disease process. Importantly, in pathological specimens where more than one member of the LOX family may be upregulated, such as in pulmonary arterial hypertension or cardiac fibrosis, the specific contributions to ECM remodeling from each of the LOX isoforms may be gleaned by selective inhibition (or knockdown/knockout) of one LOX isoform at a time. In animal studies where LOX inhibitors are delivered systemically, this in situ activity assay can be applied to identify successful LOX inhibition in the targeted organs (e.g. lung in pulmonary fibrosis) and any unintended effects in other organ systems (e.g. heart, kidney, liver in pulmonary fibrosis). Additionally, this assay can be used to verify new animal models where specific LOX proteins are knocked-down/knock-out or overexpressed.

Another potential application of this assay could be to screen new selective and specific inhibitors of each of the LOX family enzymes. This would be achieved by using cell lines that specifically express a single LOX isoform, and comparing inhibitor effects on each LOX isoform in isolation. Finally, intracellular LOXL2 activity, such as histone modifications could also be investigated with this assay. In this instance, and in cells where significant endogenous biotin is expected, a blocking step with unlabeled streptavidin before applying the BHZ would reduce background noise.

This assay has several advantages over the widely used Amplex Red HRP-coupled LOX activity assay: 1) simpler sample processing that bypasses the need for tissue homogenizing and ECM protein recovery steps; 2) broader dynamic range, evidenced by the larger activity fold change measured in A7r5 cells overexpressing LOXL2 by BHZ labeling (**Fig 2A**), compared to that measured by the traditional H_2_O_2_ coupled LOX activity assay (**Fig S3A**); 3) broader application in tissue samples, which will allow measurement of LOXs activity that correlates to in situ pathophysiological changes in animal models. Additionally, the observed assay sensitivity is higher when applied with tissue samples over cellular samples with a good signal-to-noise ratio in both cells (>4) and tissue (~10) as calculated using the grayscale intensities in specimens vs. the negative controls.

Limitations and precautions: This assay measures total LOX activity and cannot identify the activity of a single LOX. However, the contributions of specific LOX isoforms to total LOX activity in situ can be determined by the inclusion of isoform-specific inhibitors or knock-out studies. In our study, contributions from AOC3/SSAO were limited. However, in fibrotic specimens where SSAO activity can be significant, assays should be performed with and without LOX inhibitors and SSAO inhibitors to isolate the contribution of LOX to the overall BHZ signal. Another source of noise in the system could be protein carbonylation arising due to oxidative/nitrosative stress. In this instance, two additional steps and controls could be included: 1) blocking of existing carbonyls using non-biotinylated hydrazide followed by the BHZ assay; 2) inclusion of an antioxidant (e.g. N-acetyl cysteine or vitamin E) treatment to quench ROS, 3) co-immuno-fluorescence to detect colocalization of BHZ and LOXs in the samples.

In conclusion, we have developed and optimized a robust assay to detect LOX-catalyzed modification of ECM proteins to allysines. The assay distinguishes not only LOXs overexpression and depletion, as shown in our cellular studies, but also physiologically relevant activation in vascular aging. The assay is robust and reliable and is extendable to study the in situ activity of other enzymes of the LOX family.

## Materials and Methods

### Reagents

EZ-Link Hydrazide-Biotin (BHZ) was purchased from ThermoFisher. BHZ was dissolved in DMSO at the maximal soluble concentration (50 mM) for this assay. For Western blotting and immunofluorescence staining, the following antibodies were used: LOXL2 rabbit monoclonal (Abcam, ab179810), LOX rabbit polyclonal (Invitrogen, PA1-46020), LOXL1 mouse monoclonal (Santa Cruz, sc-166632), LOXL3 rabbit polyclonal (Santa Cruz, sc-68939), VAP-1 (AOC3) mouse monoclonal antibody (Santa Cruz, sc-373924), Collagen I monoclonal antibody (Invitrogen, MA1-26771), Collagen IV polyclonal antibody (Assay Biotech, C0157), GAPDH mouse monoclonal (Novus Bio, catalog #NB300221), goat anti-mouse IgG (H+L)-HRP conjugate (Biorad 1706516), AffiniPure goat anti-rabbit IgG (H+L)-HRP conjugate (Jackson ImmunoResearch, 111035144), Cy5 AffiniPure Goat Anti-Rabbit IgG (H+L) (Jackson ImmunoResearch, 111175144), Alexa Fluor 568 Goat anti-Rabbit IgG (H+L) Secondary Antibody (Invitrogen, A-11011), Alexa Fluor 488 Donkey anti-Mouse IgG (H+L) Secondary Antibody (Invitrogen, A-21202) and fluorescein (DTAF) streptavidin (Jackson ImmunoResearch, 016010084). Human LOXL2 CRISPR lentivirus was from Abm Good (K1225015). An adenovirus used to overexpress C-terminal His-tagged LOXL2 was purchased from Vector Biolabs. Vitamin E (α-Tocopherol, Sigma-Aldrich) was dissolved in ethanol to 50mM before use in cell culture. All other reagents were of the highest purity commercially available. BAPN (β-aminopropionitrile) was purchased from Sigma-Aldrich and dissolved in PBS. PAT-1251 (IUPAC name: 3-((4-(aminomethyl)-6-chloropyridin-2-yl)oxy)phenyl)((3*R*,4*R*)-3-fluoro-4-hydroxypyrrolidin-1-yl)methanone, CAS# 2098884-53-6) was obtained from Sundia Meditech Co.Ltd, and dissolved in 0.5% methyl cellulose.

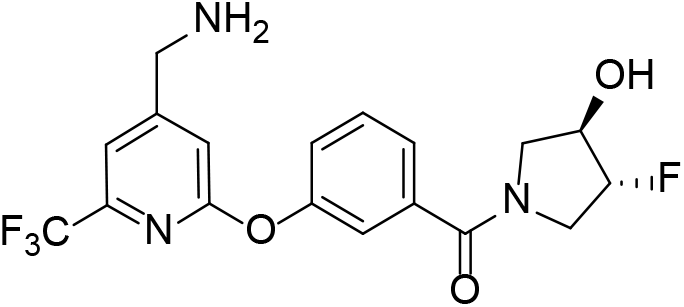

### LOXL2 cloning and mutagenesis

The LOXL2 cDNA (fusion clone) was obtained from DNAsu in pDONR entry vector. The two histidines H626/628 shown to be critical for copper binding and thus catalytic activity ^39,40^ were mutated to glutamine with the QuikChange site-directed mutagenesis kit using FP: ATC TGG CAC GAC TGT <u>CAA</u> AGG <u>CAA</u> TAC CAC AGC ATG and RP: CAT GCT GTG GTA <u>TTG</u> CCT <u>TTG</u> ACA GTC GTG CCA GAT. The resulting LOXL2 H626/628Q double mutant (LOXL2-DM) was confirmed by Sanger sequencing. The LOXL2-DM was then transferred to pAd-DEST (ThermoFisher) and used to generate adenovirus Ad-LOXL2-DM.

### Cells

A7r5 (ATCC) rat thoracic artery smooth muscle cells were sub-cultured in Dulbecco’s modified Eagle media (DMEM; Gibco) supplemented with 10% fetal bovine serum and antibiotic-antimycotic (Gibco). HASMCs (ThermoFisher) were cultured in smooth muscle cell media (ScienCell) containing 2% fetal bovine serum, SMC-growth supplement, and antibiotic-antimycotic. As previously described^12^, LOXL2 gene was targeted by using a human LOXL2 lentivirus set containing three gRNAs — T1 (TGTACTATGATGGCCAG), T2 (GCTTGCGGTAGGTTGAG), and T3 (ATGTCACCTGCGAGAAT) — followed by selection with puromycin (2.5 μg/mL; ThermoFisher). The T1 gRNA efficiently disrupted LOXL2 expression and was used to generate LOXL2-depleted HASMC T1 cells in these studies.

### Mouse model

LOXL2+/− heterozygous mice were produced by using cryopreserved sperm in which the LOXL2 gene was disrupted by insertion of a long terminal repeat-trapping cassette into exon 1 (Taconic Inc.). The mice were bred in-house and genotyped as previously described^12^. Male LOXL2+/− and WT littermate mice were used in this study. Animals were maintained in the AAALAC accredited Johns Hopkins University vivarium with a 12-h light/dark cycle and were fed and watered ad libitum. All animal protocols were approved by the Johns Hopkins Institutional Animal Care and Use Committee.

### Western blotting

Aorta samples were homogenized in RIPA buffer supplemented with protease inhibitor cocktail (Roche). Soluble proteins and insoluble matrix were separated by centrifugation at 10,000 × *g*. The insoluble matrix was then resuspended in reducing 2% (w/v) SDS lysis buffer and boiled for 15 min. Samples were fractionated by SDS-PAGE and transferred onto nitrocellulose membrane by electrophoresis. Membranes were blocked in 3% nonfat milk in TBST (Tris-buffered saline, 0.1% Tween 20) and then incubated with primary antibody (1:1000, 2 h) and secondary antibody (1:10,000, 2 h). Membranes were washed in TBST and developed with the Clarity Western ECL system (Bio-Rad).

### In vitro LOXL2 BHZ binding assay

A7r5 cells were transduced with adenovirus in serum-free insulin-transferrin-selenium (ITS)-supplemented DMEM for 48 h to overexpress LOXL2 or LOXL2-DM. Cell culture medium containing overexpressed LOXL2 or LOXL2 DM was collected and incubated with increasing concentrations of BHZ (0, 50, 100, 200, 300 μM) at 4°C for 2 h or 24 h with gentle rocking. Biotinylated proteins were enriched by using Pierce Streptavidin Agarose beads (Thermo Scientific) according to the manufacturer’s protocol. Beads were then washed three times in PBS, and bound proteins were recovered by boiling in reducing Laemmli buffer. Samples were analyzed by Western blotting.

### Cellular in situ LOXL2 activity assay

Cells were seeded on coverslips at 80% confluence, allowed to adhere, and serum-starved overnight. To induce overexpression of LOXL2 or LOXL2-DM, cells were transduced with adenovirus for 24 h before initiating the activity assay. BHZ (0-150 μM) was then delivered to confluent monolayers of cells for 0–72 h in ITS-supplemented DMEM (serum-free). Samples were then rinsed free of excess BHZ twice with sterile PBS and fixed in 3.7% formaldehyde for 30 minutes before proceeding to fluorescence staining.

### Aortic tissue LOXL2 activity assay

Aortas from young (<3 months old) and old (>18 months old) WT and LOXL2+/− littermates were extracted and cleaned free of connective tissue in Krebs-HEPES buffer containing 0.25 μg/mL fungizone (Gibco). The aortas were then cut into 5 mm rings and transected open. Rings were incubated in DMEM + ITS at 37°C with or without 200 μM BHZ for 24 h. After excess BHZ was rinsed away with sterile PBS (3 times, 15 min each), the samples were fixed in 3.7% formaldehyde for 1 h before immunofluorescence staining. For imaging, the aortic rings were pinned en face with endothelium facing up on silastic-coated dishes and imaged by upright confocal microscopy (Zeiss 710NLO). Z-stack images were taken for the thickness of 50–100 μm and the maximum intensity projection on the 3-dimensional images along the Z-axis was obtained to capture the signal from the entire thickness of the aorta. LOX, LOXL1, LOXL2, LOXL3, AOC3 expression in aortic tissue was verified by Western blotting.

### Fluorescence staining

Fixed samples were blocked in 3% bovine serum albumin in PBS for 1 h. LOXs and AOC3 were labeled by incubating the samples with primary antibody (1:150, 2 h) followed by fluorescein-conjugated secondary antibody (1:250, 2 h). Biotinylated allysine hydrazones were detected by fluorescein (DTAF) streptavidin (1:250, 2 h). Nuclei were labeled with DAPI (1 μg/mL, 15 min). All staining procedures were performed at room temperature. Samples were imaged by confocal microscopy (Zeiss LSM700) and at least 3 images were obtained for each specimen. Fluorescence intensity was measured based on mean gray value (Image J). Fluorescence intensity measured in negative controls (samples without BHZ incubation) was averaged and subtracted from sample measurements to adjust for background in the bar graphs.

### Fluorometric lysyl oxidase assay

Conditioned cell culture medium was collected and concentrated 20-fold by using Amicon Ultra-0.5 mL 10K MWCO Centrifugal Filters (Millipore Sigma). LOX activity was measured with an Amplite Fluorimetric Lysyl Oxidase Assay kit (AAT Bioquest) according to the manufacturer’s protocol. Briefly, 50 μL of concentrated conditioned cell culture medium was added to 50 μL of LOX working solution containing LOX substrate, horseradish peroxidase, and Amplite horseradish peroxidase substrate and incubated at 37°C for 1 h, during which fluorescence intensity was measured at Ex/Em = 540/590 nm by a fluorescence spectrophotometer. H_2_O_2_ released by the LOX oxidation was detected in the HRP-Amplex Red coupled reaction. LOX reaction rate was calculated as the slope of fluorescence increase over time.

